# CTP Drives Human CTPS1 Filamentation

**DOI:** 10.1101/2025.02.22.639624

**Authors:** Chen-Jun Guo, Xiaojie Bao, Ji-Long Liu

**Author notes:** Correspondence: Ji-Long Liu. These authors contributed equally: Chen-Jun Guo and Xiaojie Bao.

## Abstract

CTP synthase (CTPS) is a key enzyme in *de novo* CTP synthesis, playing a critical role in nucleotide metabolism and cellular proliferation. Human CTPS1 (hCTPS1), one of the two CTPS isoforms, is essential for immune responses and is highly expressed in proliferating cells, making it a promising therapeutic target for immune-related diseases and cancer. Despite its importance, the regulatory mechanisms governing hCTPS1 activity remain poorly understood. Here, we reveal that CTP, the product of CTPS, acts as a key regulator for hCTPS1 filamentation. Using cryo-electron microscopy (cryo-EM), we resolve the high-resolution structure of CTP-bound hCTPS1 filaments, uncovering the molecular details of CTP binding and its role in filament assembly. Importantly, we demonstrate that CTP generated from the enzymatic reaction does not trigger filament disassembly, suggesting a conserved regulatory pattern. Furthermore, by analyzing the binding modes of two distinct CTP-binding pockets, we provide evidence that this filamentation mechanism is evolutionarily conserved across species, particularly in eukaryotic CTPS. Our findings not only elucidate a novel regulatory mechanism of hCTPS1 activity but also deepen the understanding of how metabolic enzymes utilize filamentation as a conserved strategy for functional regulation. This study opens new avenues for targeting hCTPS1 in therapeutic interventions.

## INTRODUCTION

Cytidine triphosphate synthase (CTPS) is a pivotal metabolic enzyme that sits at the crossroads of nucleotide biosynthesis, playing a central role in the synthesis of DNA, RNA, and phospholipids(1, 2, 3, 4, 5). As the rate-limiting enzyme in the *de novo* CTP synthesis pathway, CTPS not only drives cellular proliferation but also tightly regulates the intracellular CTP pool, making it essential for maintaining metabolic homeostasis(6, 7, 8, 9, 10).

Structurally, CTPS functions as a homologous tetramer, with each monomer comprising an N-terminal amidoligase (AL) domain and a glutamine amidotransferase (GAT) domain(11, 12). The enzyme catalyzes a multi-step reaction: the AL domain phosphorylates uridine triphosphate (UTP) using ATP, generating an unstable intermediate, 4-phospho-UTP (4-Pi-UTP)(13, 14). This intermediate is then aminated by ammonia, which is hydrolyzed from glutamine in the GAT domain and transported through an intramolecular ammonia tunnel(14, 15). The reaction is finely tuned by allosteric regulators such as GTP, which stabilizes intermediate states, and feedback-inhibited by the end product, CTP(14, 16, 17, 18, 19). These regulatory mechanisms, along with the tetrameric structure, are highly conserved across species, underscoring the fundamental role of CTPS in cellular metabolism(11, 20, 21, 22, 23).

Beyond its enzymatic activity, CTPS exhibits a remarkable ability to polymerize into large-scale filamentous structures known as cytoophidia(24). These structures have been observed in diverse organisms, including bacteria, yeast, fruit flies, and human cells, suggesting an evolutionarily conserved regulatory mechanism(24, 25, 26, 27, 28). Filamentation is thought to modulate CTPS activity, although the precise mechanisms remain elusive(14, 24, 25, 26, 27, 28, 29, 30).

In several mammals, including humans, CTPS exists as two isoforms—CTPS1 and CTPS2—with 75% sequence identity but distinct physiological roles(7). Human CTPS1 (hCTPS1) is widely expressed in proliferating cells and is critical for immune function, as its deficiency leads to severe immune impairment(6, 7). Notably, CTPS1 is overexpressed in many cancer cells, and its activity is essential for the replication of viruses, pathogens, and parasites, making it a promising therapeutic target for cancer, infectious diseases, and immune disorders(4, 31, 32, 33, 34, 35, 36, 37, 38, 39). Despite its biomedical significance, the regulatory mechanisms governing hCTPS1 activity, particularly its filamentation dynamics, remain poorly understood.

Previous studies on eukaryotic CTPS, including Drosophila CTPS (DmCTPS) and hCTPS2, have revealed that filamentous structures adopt two distinct conformations, enabling enzymatic regulation through ligand binding and release(23, 29). For hCTPS1, it has been proposed that filament formation is induced by substrate binding, while product binding (CTP) triggers filament disassembly(30). However, due to limitations in structural resolution and experimental conditions, the molecular details of how CTP binding regulates hCTPS1 filamentation and enzymatic activity remain unclear.

In this study, we address this critical gap by investigating the impact of CTP on hCTPS1 filamentation. Using cryo-electron microscopy (cryo-EM), we resolved the high-resolution (3.3 Å) structure of hCTPS1 filaments bound to CTP, revealing the precise binding modes and conformational changes associated with filament assembly. Our findings demonstrate that CTP, the enzymatic product, does not induce filament disassembly, challenging the prevailing model of hCTPS1 regulation. Furthermore, we provide evidence that this filamentation mechanism is evolutionarily conserved, particularly in eukaryotic CTPS. These insights not only elucidate a novel regulatory mechanism of hCTPS1 but also deepen our understanding of how metabolic enzymes utilize filamentation as a conserved strategy for functional regulation. This work opens new avenues for targeting hCTPS1 in therapeutic interventions for cancer, immune disorders, and infectious diseases.

## RESULTS

### hCTPS1 assembles into large filamentous structures

hCTPS1 functions as a homologous tetramer, with each monomer comprising a glutamine amidotransferase (GAT) domain and an amidoligase (AL) domain. In the presence of GTP, the GAT domain hydrolyzes glutamine (Gln) to produce ammonia, which is transferred to the AL domain to catalyze the conversion of uridine triphosphate (UTP) to cytidine triphosphate (CTP) via an ATP-dependent intermediate, 4-phospho-UTP (4-Pi-UTP) (Figure 1A). Purified hCTPS1 exhibited high homogeneity and robust catalytic activity in enzyme assays (Figure 1B).

**Figure 1.**
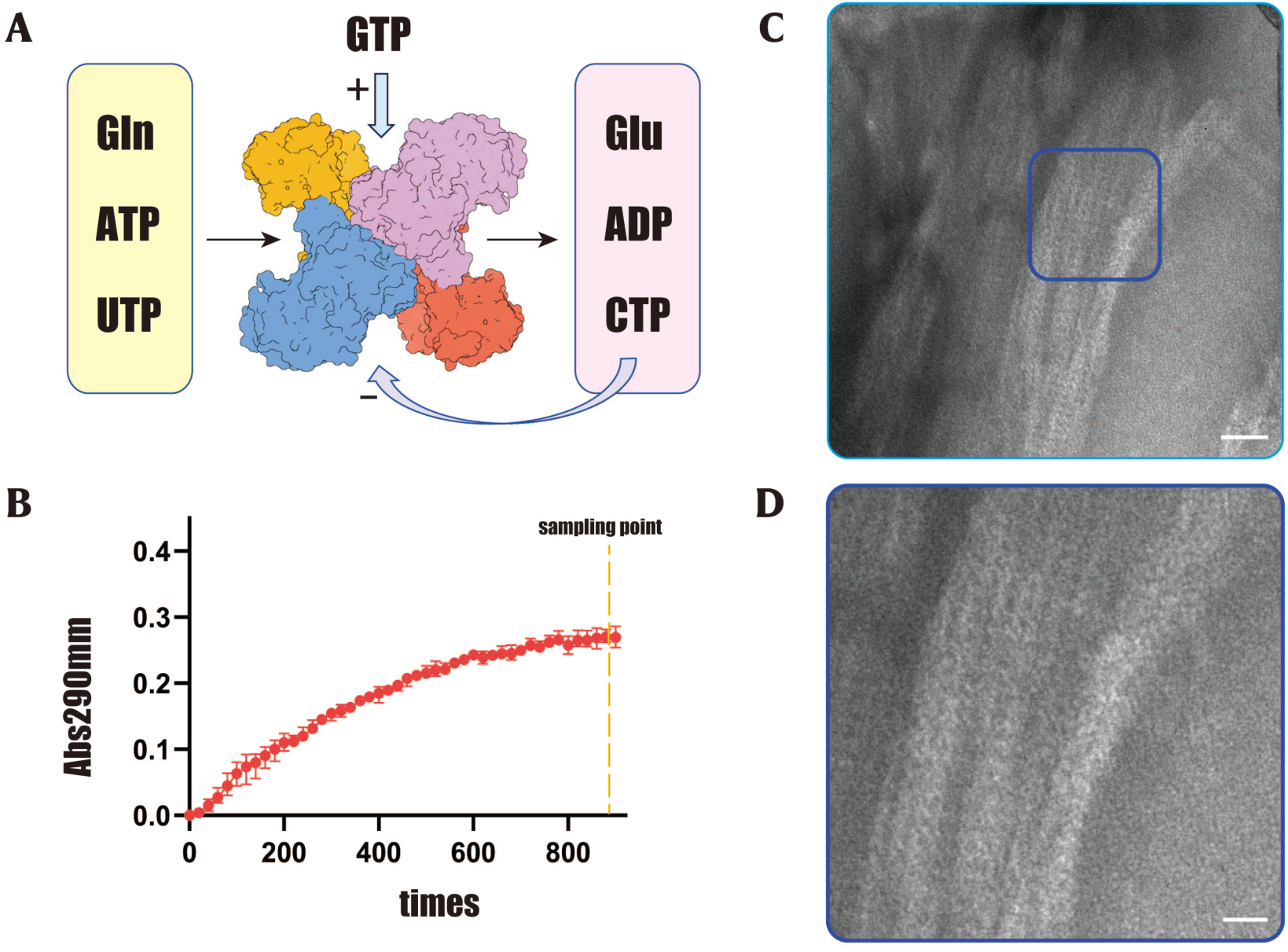
Catalytic mechanism and enzymatic reaction of hCTPS1. The way of forming CTP reactions catalyzed by CTP synthase. UTP accepts Phosphoric acid hydrolyzed by ATP and form an intermediate 4-Pi-UTP in GAT domain (grey box). The ammonia is released from glutamine by the allosteric regulator GTP in AL domain (yellow box) and then ligated to 4-Pi-UTP to form CTP in GAT domain. (A) Kinetic curves of hCTPS1 enzyme assay in 0.4 mM GTP and none GTP system. The X-asis indicates the increase of time. The Y-axis indicates the change in absorbance of the enzymatic reaction. (C-D) Negative-staining EM micrographs of hCTPS1 at the end of enzymatic reaction. The scale bar is 50 nm. (D) is the zoomed-in view of the top yellow box in (C). The scale bar is 200 nm.

In the enzymatic reaction system containing magnesium ions, GTP, and excess Gln, the reaction produced ADP, glutamate (Glu), and CTP. After 1200 seconds of reaction with 1 mM UTP and 1 mM ATP, the system generated approximately 0.45 mM CTP, achieving a conversion rate of 45% and a final UTP:CTP ratio of 11:9. Strikingly, cryo-electron microscopy (cryo-EM) analysis of the reaction solution revealed large, stacked bundles composed of periodically arranged filamentous structures (Figure 1C, D). Grayscale analysis confirmed the uniformity of these structures, suggesting that hCTPS1 has the ability to form filaments and further aggregate in CTP-rich environments.

### CTP induces hCTPS1 filamentation

To investigate the mechanism underlying hCTPS1 filamentation, we explored the effects of various ligands on its self-assembly. Previous studies have shown that hCTPS1 forms filaments in the presence of substrates (ATP and UTP) and the allosteric regulator GTP (Figure 2A). Here, we demonstrated that hCTPS1 also forms filaments in solutions containing the enzymatic products CTP, ADP, and Glu (Figure 2B), indicating that product binding can induce filament assembly.

**Figure 2.**
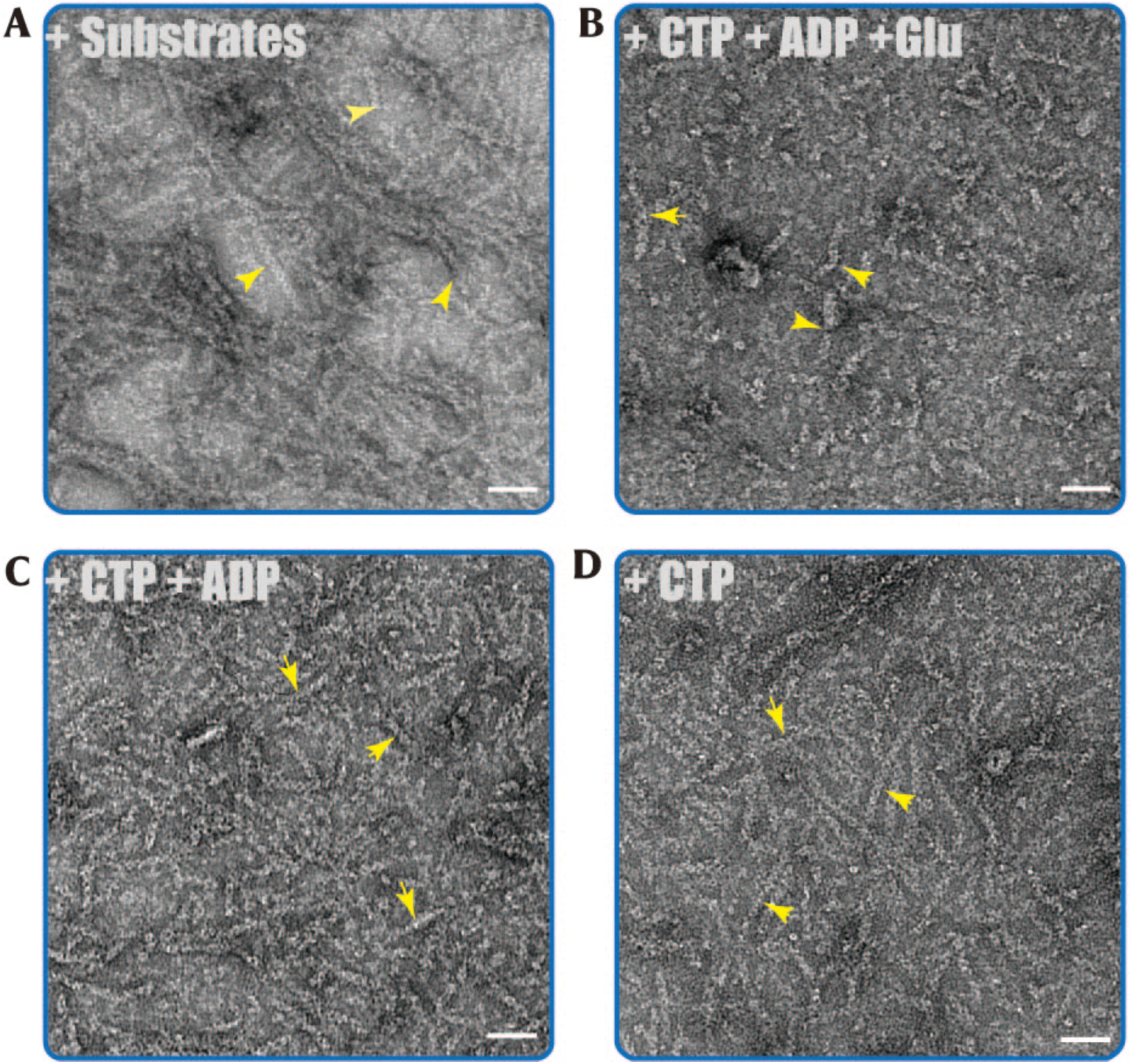
Negative staining of human hCTPS1 filament. The wild-type human hCTPS1 can form filament under several condition with Mg^2+^. The scale bar is 50 nm. Filaments are indicated by yellow arrows. (A) Negative-staining EM micrographs of hCTPS1 incubated with whole substrates (ATP, UTP, GTP and Gln) combinations. (B) Negative staining of hCTPS1 incubated under whole product ligands condition. (C) A solution system containing only CTP and ADP can form hCTPS1 filamentation. (D) hCTPS1 binding with single ligand CTP can be induced to polymerize.

By systematically reducing the types of ligands in the reaction system, we identified CTP and ADP as key drivers of hCTPS1 filamentation. Remarkably, hCTPS1 formed filaments in the presence of CTP alone (Figure 2C) or ADP alone, but not in systems containing only Glu. These results suggest that CTP, the enzymatic product, can coexist with hCTPS1 filaments, challenging the prevailing view that CTP triggers filament disassembly.

To further validate this finding, we conducted a time-resolved experiment in which CTP was added to hCTPS1 and immediately imaged using negative-staining electron microscopy. Filamentous structures were observed within seconds, confirming that CTP binding rapidly induces hCTPS1 filamentation (Figure S1). This finding contradicts previous models and suggests that CTP-mediated feedback inhibition of hCTPS1 does not involve filament disassembly.

### Cryo-EM structure of CTP-bound hCTPS1 filaments

To elucidate the structural basis of hCTPS1 filamentation under product-bound conditions, we resolved the cryo-EM structure of hCTPS1 filaments in the presence of CTP (Figure S2&S3). To stabilize the GAT domain, we included 6-diazo-5-oxo-L-norleucine (DON), a glutamine analog that irreversibly inhibits CTPS activity by covalently binding to the glutamine site. Negative-staining experiments confirmed that DON does not disrupt filament formation.

Cryo-EM analysis revealed clear filamentous structures, and 2D classification highlighted detailed structural features (Figure 3A). Using 312,573 particles for reconstruction, we achieved a final resolution of 3.3 Å. The structure showed that hCTPS1 filaments consist of helical arrays of tetramers, with each helical unit rising by 105 Å and twisting by approximately 36° (Figure 3B, C). The tetramers are connected through α-helix-mediated interactions involving residues 346–357 in the GAT domain. Notably, residues H355 and W358, which are critical for filament assembly across species, exhibited continuous electron density in the density map, underscoring their conserved role (Figure 3D). Each tetramer contained two distinct CTP binding sites, providing insights into the molecular basis of CTP-mediated regulation.

**Figure 3.**
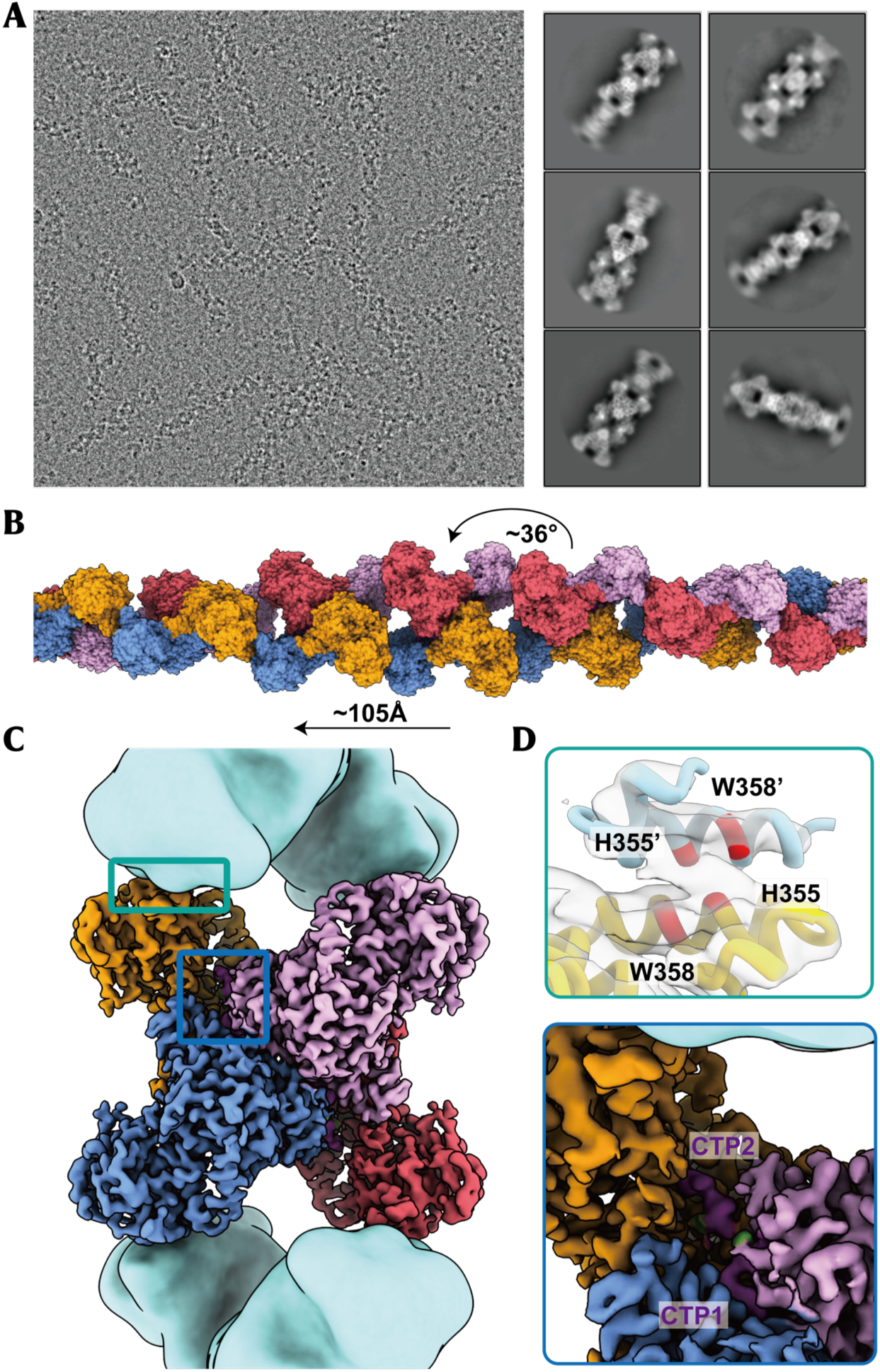
Cryo-electron microscopy (cryo-EM) analysis and overrall structure of hCTPS1 filaments with CTP. (A) Cryo-electron micrograph of hCTPS1 under CTP and Don condition. Several representative 2D class averages of the hCTPS1 filament are selected in view of the relion4 on the right. (B) Reconstructed model of hCTPS1 filament. CTP bound filament model twists 36 ° between two adjacent tetramers and shifts along the axis 105Å (C) Cryo-EM structure of hCTPS1 bound to CTP and Don. The central hCTPS1 tetramer is colored by different protomers. (D) Zoomed-in view of the top green box in (C), showing the interface of two adjacent tetremers. The density of residues responsible H355 and W358 for the interactions are color red. Zoomed-in view of the below blue box in (C), showing the CTP in the non-canonical site.

### Conserved CTP binding modes in hCTPS1 filaments

CTPS is known to undergo product feedback inhibition, where high CTP concentrations induce conformational changes that reduce enzymatic activity. In our structure, two CTP molecules were observed bound to each protomer: one at the interface of three protomers (overlapping with the UTP binding site) and the other within a single protomer (overlapping with the ATP binding site) (Figure 4A). We designated these as the canonical and non-canonical binding pockets, respectively.

**Figure 4.**
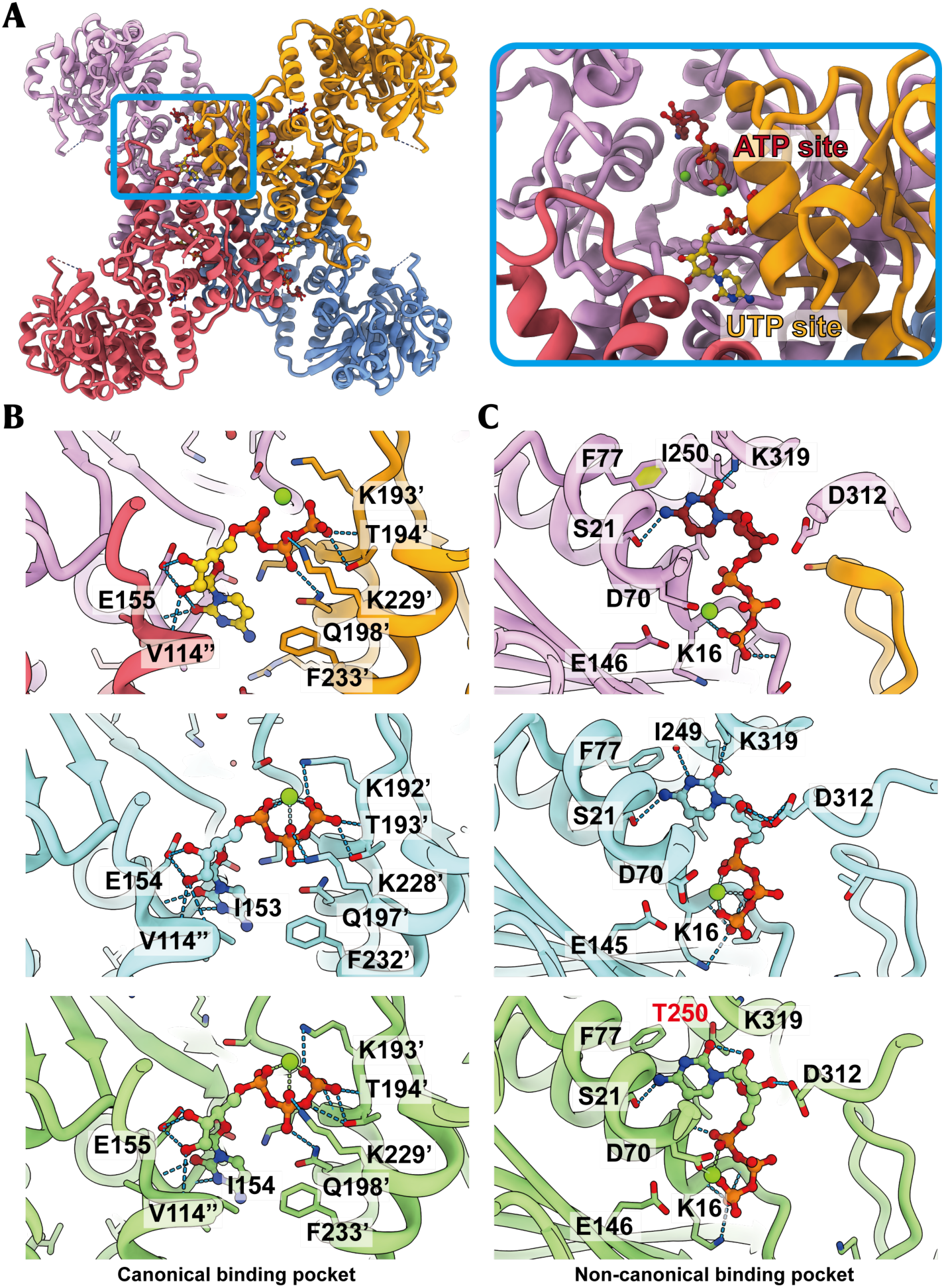
CTP binding modes in hCTPS1 and comparison of CTP binding pockets among organisms. (A) A single refined hCTPS1 tetramer model from the filament, colored by protomer (pink, goden, blue and red) and nucleotide highlighted in yellow and crimson. Zoomed-in view of the below blue box shows the CTP in the canonical(yellow) and non-canonical(crimson) site, which are respectively occupied by UTP and ATP under substrate state. (B-C) Diversities of CTP binding pockets among organisms. Models of hCTPS1, Drosophila cytidine triphosphate synthase(DmCTPS) and hCTPS2 are colored in pink, blue, and green, respectively, with the corresponding Protein Data Bank (PDB) codes 9lyx, 7dpw, and 7mh1. (B) Canonical CTP binding pocket. CTP is represented by yellow sphere-and-stick models, hydrogen bonds are shown as blue dashed lines, and magnesium ions(Mg) represented by green spheres. (C) Comparisons of non-canonical CTP binding sites.

In the canonical binding pocket, the triphosphate group of CTP is stabilized by electrostatic and hydrogen-bonding interactions with protomer No. 3. Residue E155 forms salt bridges with the ribose group, while the cytosine base interacts electrostatically with the backbone of E155. Sequence alignment revealed that the residues involved in CTP recognition in this pocket are highly conserved in hCTPS2 and Drosophila CTPS (DmCTPS) (Figure 4B & Figure S5, 6).

In the non-canonical binding pocket, the triphosphate group interacts electrostatically with a loop, while the cytosine group is stabilized by interactions with S21, K319, and F77. These residues are identical in DmCTPS and highly conserved in hCTPS2, except for T250, which differs between hCTPS1 (I250) and hCTPS2 (T250). This difference has been exploited to design selective inhibitors targeting hCTPS1 (Figure 4C)(40).

### Conserved conformational changes in hCTPS1 filaments

Like hCTPS2 and DmCTPS, hCTPS1 filaments exhibit distinct conformations under substrate-bound and product-bound conditions. While the filament assembly interface remains unchanged, the helical parameters of the filaments vary with ligand binding (Figure 5A & Figure S7).

**Figure 5.**
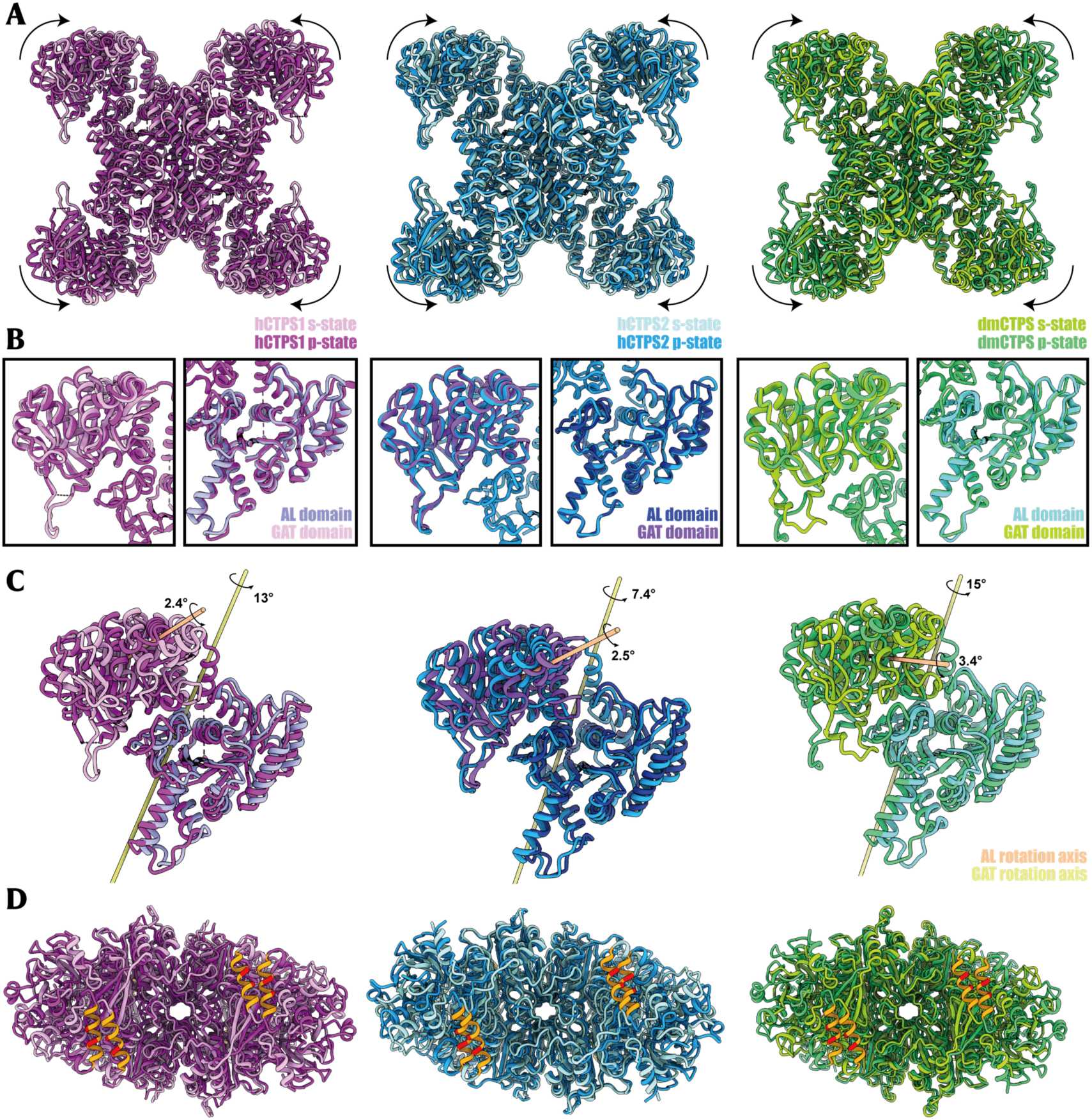
Structural comparison between hCTPS1(purple), hCTPS2(blue) and DmCTPS(green) in substrate-state and product-state. The structural of hCTPS1, hCTPS2 and DmCTPS in s-state are colored in pink, powder blue and lime green, with the corresponding Protein Data Bank (PDB) codes 7mgz, 6pk4, and 7dpt. The structural of hCTPS1, hCTPS2 and DmCTPS in p-state are colored in purple, deep sky blue and forest green, with the corresponding Protein Data Bank (PDB) codes 9lyx, 7mh1, and 7dpw. (A) Structural comparison between hCTPS1 tetramer, hCTPS2 tetramer and DmCTPS tetramer in substrate-state and product-state. (B) Structural comparison of AL domain and GAT domain. The wing structure in GAT domain has deviated among the three species, while the conformation of AL domain remains almost unchanged. (C) Rotation axis of AL domain and GAT domain transitioning between two states. (D) Comparison of the interaction interface between adjacent tetramers in filament. The responsible H355 and W358 for the interactions are color red.

Structural alignment of the AL domain (residues 1–270) between substrate-bound (s-state) and product-bound (p-state) hCTPS1 revealed an RMSD of 1.267 Å, with conformational differences localized to the Arch region (residues 45–67) and Loop183–194. The GAT domain exhibited high rigidity, with an RMSD of 0.899 Å for all 249 residue pairs, indicating stable conformations under different ligand conditions (Figure 5B).

Further analysis showed that the conformational changes in the hCTPS1 tetramer primarily involve rotational displacement of the GAT domain, while the AL domain remains relatively fixed (Figure 5C). The filament interface on the GAT domain surface maintains a stable, outward-facing conformation, enabling antiparallel assembly of tetramers. This assembly mode is conserved in hCTPS2 and DmCTPS (Figure 5D).

In summary, hCTPS1 filaments exhibit conserved conformational changes and assembly interfaces under substrate and product conditions, highlighting the evolutionary conservation of the filamentation regulatory pattern.

## DISCUSSION

### CTP promotes hCTPS1 filamentation

Previous studies suggested that hCTPS1 filaments are stable only in the presence of substrates, and that binding to the product CTP triggers filament disassembly, leading to negative feedback inhibition(30). However, our findings challenge this model by demonstrating that hCTPS1 can form stable filaments in the presence of CTP. We resolved the high-resolution structure of CTP-bound hCTPS1 filaments, revealing that CTP generated during enzymatic reactions does not induce filament depolymerization. This discovery provides a new perspective on the regulatory mechanisms of hCTPS1 activity.

The two CTP binding pockets in hCTPS1 exhibit ligand recognition modes highly similar to those of Drosophila CTPS (DmCTPS) and hCTPS2(14, 29). The canonical binding pocket, located at the interface of three monomers, overlaps with the UTP binding site, while the non-canonical binding pocket occupies the ATP/ADP binding site. Although the competitive binding mechanism at the non-canonical site remains to be fully elucidated, our structural data show that the phosphate backbone of CTP shares binding pockets with UTP and ATP, highlighting a conserved mechanism of ligand recognition and stabilization across species.

Notably, the formation of hCTPS1 filaments in the presence of CTP is rapid and robust. Filament assembly occurs almost immediately after incubation, and prolonged incubation leads to the formation of large, stable bundles. This suggests that the filamentous structure of hCTPS1 bound to CTP is highly stable, further supporting the idea that CTP-mediated regulation does not involve filament disassembly.

### Filamentation of hCTPS1 and hCTPS2

hCTPS1 and hCTPS2 play critical roles in cell proliferation and immune function, yet their precise molecular mechanisms remain poorly understood(6, 7). hCTPS1, in particular, is essential for the proliferation of cancer cells, such as Jurkat cells, and its inhibition is considered a promising therapeutic strategy for cancer treatment(4, 7, 31, 32, 33, 37, 38, 39). Additionally, hCTPS1 is implicated in severe human diseases and post-transplant medication, making it a valuable drug target. A detailed understanding of its ligand binding and filamentation mechanisms is crucial for developing hCTPS1-specific inhibitors.

Our study reveals that the filament interfaces of hCTPS1, hCTPS2, and DmCTPS are highly conserved, with interactions mediated by an α-helix (residues 346–357). These interfaces differ from those of *Escherichia coli* CTPS (EcCTPS), highlighting evolutionary divergence between prokaryotic and eukaryotic CTPS. Despite these differences, the rotational displacement of the GAT and AL domains during ligand binding and release is conserved, suggesting a shared regulatory mechanism.

This study addresses a critical gap in our understanding of hCTPS1 filamentation. While the conformational changes observed in CTP-bound filaments do not fully explain the transition between filaments and tetramers, they provide valuable insights into the regulation of hCTPS1 activity. The discovery of the CTP-bound hCTPS1 filament structure offers a reliable foundation for the development of targeted therapies.

### CTPS filamentation in eukaryotes and prokaryotes

Filamentation is a conserved regulatory mechanism among metabolic enzymes, influenced by ligand binding, nutrient availability, and pH. In eukaryotes, DmCTPS and hCTPS2 can form filaments under both substrate-bound and product-bound conditions, with substrate-bound filaments exhibiting enhanced enzymatic activity. In contrast, EcCTPS forms filaments only in the presence of products, which inhibit enzyme activity.

We propose that metabolic enzymes, including hCTPS1, regulate complex physiological functions through conformational changes between free polymers and filaments. These structural changes enable efficient ligand binding and release, facilitating enzymatic reactions and maintaining dynamic equilibrium. In vivo, hCTPS likely assembles from monomers into tetramers, which then form filaments and bundle into cytoophidia. These structures persist throughout the cell cycle, rapidly regulating enzyme activity and other biochemical functions.

Despite the high sequence identity (∼75%) between hCTPS1 and hCTPS2, their filament assembly and interfaces are highly conserved, with only minor differences in helical parameters. The relative independence of the GAT and AL domains during conformational changes suggests that hCTPS filaments *in vivo* may not consist solely of homologous monomers (Figure S8). Future studies should explore the assembly mechanisms and functions of hCTPS filaments and cytoophidia in their native cellular context.

## Data availability

The structure data accession codes are EMD-63516 and PDB-9LYX.

## Acknowledgments

We thank Chia-Chun Chang for providing the plasmid. We thank Yi-Lan Li and Jia-Li Lu for technical support. EM data were collected at Bio-Electron Microscopy Facility of ShanghaiTech University. We thank the Molecular and Cell Biology Core Facility (MCBCF) at the School of Life Science and Technology, ShanghaiTech University, Bio-Electron Microscopy Facility of ShanghaiTech University and Shanghai Frontiers Science Center for Biomacromolecules and Precision Medicine for providing technical support. This work was supported by grants from: Ministry of Science and Technology of China (No. 2021YFA0804700), National Natural Science Foundation of China (Grant Nos. 32370744 and 32350710195), and UK Medical Research Council (grant nos. MC_UU_12021/3 and MC_U137788471) for grants to J.L.L.

## Author contributions statement

C.J.G and J.L.L initiated the project. X.B performed protein expression, purification and the functional assays. C.J.G and X.B prepared samples for EM studies, collected cryo-EM data, the cryo-EM data processing, model building, structure refinement, analyzed experiments, visualized results and wrote the initial manuscript.

J.L.L revised manuscript, obtained the funding and supervised the project.

## METHODS

### Purification of hCTPS1

hCTPS1 was expressed in Saccharomyces cerevisiae strain BY4741, as given by Chia-Chun Chang and Yi-Lan Li of Liu Lab, which directs expression of C-terminal 6× His-tagged hCTPS1 from the GAL1 promoter. BY4741 were grown in 1× YPD media and induced in 1× YPG media at 30℃. Cells were collected and pelleted by centrifugation at 4 ℃ at 4,000 rpm for 10 min. Cell pellets were resuspended in lysis buffer (500 mM NaCl, 50 mM Tris-HCl, 10% glycerol, 1mM PMSF, 5mM β-mercaptoethanol, 5mM benzamidine, 1 μg/mL leupeptin, pH8.0). Lysates were clarified by centigugation at 10,000 g for 45 min at 4 ℃ in a Beckman Avanti JXN-26 JA-25.50 rotor. Supernatant was incubated with equilibrated Ni-Agarose (Qiagen, Germany) for 1 h. The column was eluted with lysis buffer with 250 mM imidazole and 1 mM β-Me at pH8.0. Elution was concentrated, flash-frozen in liquid nitrogen and stored at-80℃.

### Negative-stain electron microscopy

5 μM hCTPS1 protein with diverse ligands was dissolved in a buffer containing 25 mM Tris HCl, 150 mM NaCl, pH8.0 and 10 mM MgCl2, 10 mM DTT to form filaments. Samples for negative-stain EM were prepared by loading 5.8 μl hCTPS1 to hydrophilic carbon-coated grids (400mech, Zhongjingkeyi Technology Co., China) and staining with 0.1% uranyl formate. Solution systems were incubated alternately at 23.5 ℃ and zero for 10 min, each temperature sustaining 2-5 min, or at 23.5 ℃ for 15-20 min continuously. Imaging was on a 120 kV microscope (Talos L120C, ThermoFisher, USA) with an Eagle 4 K× 4 K CCD camera system (Ceta CMOS, ThermoFisher, USA) and images were acquired at 57,000× magnification.

### Cryo-EM sample preparation and data collection

To prepare samples for cryo-EM, hCTPS1 was applied to glow-discharge amorphous alloy film (No. M01-Au300-R1.2/1.3) and blotted 2.7-3 μl, then plunged into liquid ethane using a FEI Vitrobot (4 ℃ temperature, 3-3.5 s blotting time,-1 to 1 blot force). For preparing the CTP-binding hCTPS1 sample, 5 μM hCTPS1 protein was incubated in the buffer with 2 mM CTP, 0.6 μM DON, 25 mM Tris HCl and 150 mM NaCl. Solution systems were incubated alternately at 23.5 ℃ and zero for 10 min, each temperature sustaining 2-5 min, or at 23.5 ℃ for 15-20 min continuously. Each sample is diluted twice before boltting. Images were taken with a Gatan K3 summit camera on a FEI Titan Krios electron microscope operated at 300 kV. The magnification was 29,000× in super-resolution mode with the defocus rage −1.5 to −2.5 μm and a pixel size of 0.41 Å. The total dose was 60 electrons/Å^2^ subdivided into 40 frames at 2.4-s exposure using SerialEM.

### Cryo-EM Data Processing

The whole workflow was done in RELION GUI. Moview were dose-weighted, aligned and summed by MotionCor2 through Relion. CTF parameters were assessed by CTFFIND4. Respectively, 95,054 particles were automatically picked and performed 2-dimensional (2D) classification. After 2-dimensional and 3-dimensional (3D) classification, 62,319 particles were selected for the 3D refinement to generate two maps of 3.5Å. CTF refinement and Bayesian polishing were applied to each particle. Finally, we constructed maps with resolution 3.3 Å.

### Model Building and Refinement

Human CTPS1 monomer model from AlphaFold was applied for the initial models. The tetramer models were generated and docked into the electron density map by Chimera v.1.14. Manual adjustment and rebuilding were performed with Web-coot(41). Phenix was used to achieve real space refinements. All figures were generated using UCSF Chimera, ChimeraX and Phenix(42, 43).

### hCTPS1 enzyme assay

8μM purified hCTPS1 at concentrations as described in each experiment was incubated in the buffer containing 150mM NaCl, 25mM Tris-HCl pH 8.0, 1 mM ATP, 1 mM UTP, 0.4 mM GTP, and 10 mM MgCl2 for 20 min at 37℃. 10mM Gln was added to the mixture after prewarmed as initiator. Absorbance at 291 nm of each mixture(150ul) was measure using SpectraMax i3 to indicate the production of CTP, and measurements were made at 20s intervals for at less 30min until the absorption value tended to be stable(44).

## Supplementary info

**Supplementary Tables S1**

**Supplementary Figures S1-S8**

**Figure S1.**
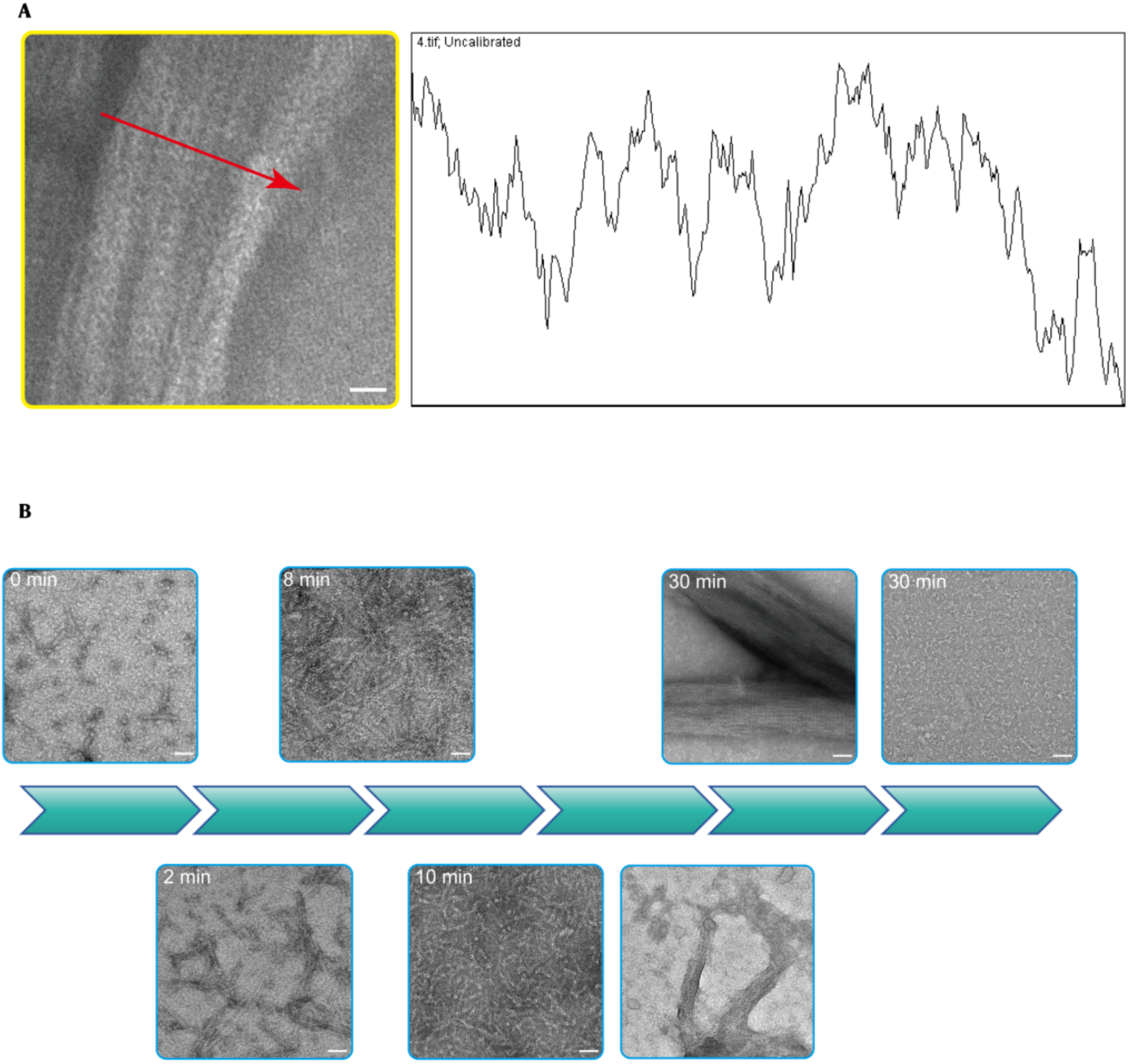
Grayscale analysis of bundle structure and time experiment of hCTPS1 combined with CTP. (A) The grayscale map shows the density information of the bundle structure from left to right for the area pointed by the red arrow. (B) Negative-staining EM micrographs of hCTPS1 binding with CTP at 0, 2,8 10, 30 min.

**Figure S2.**
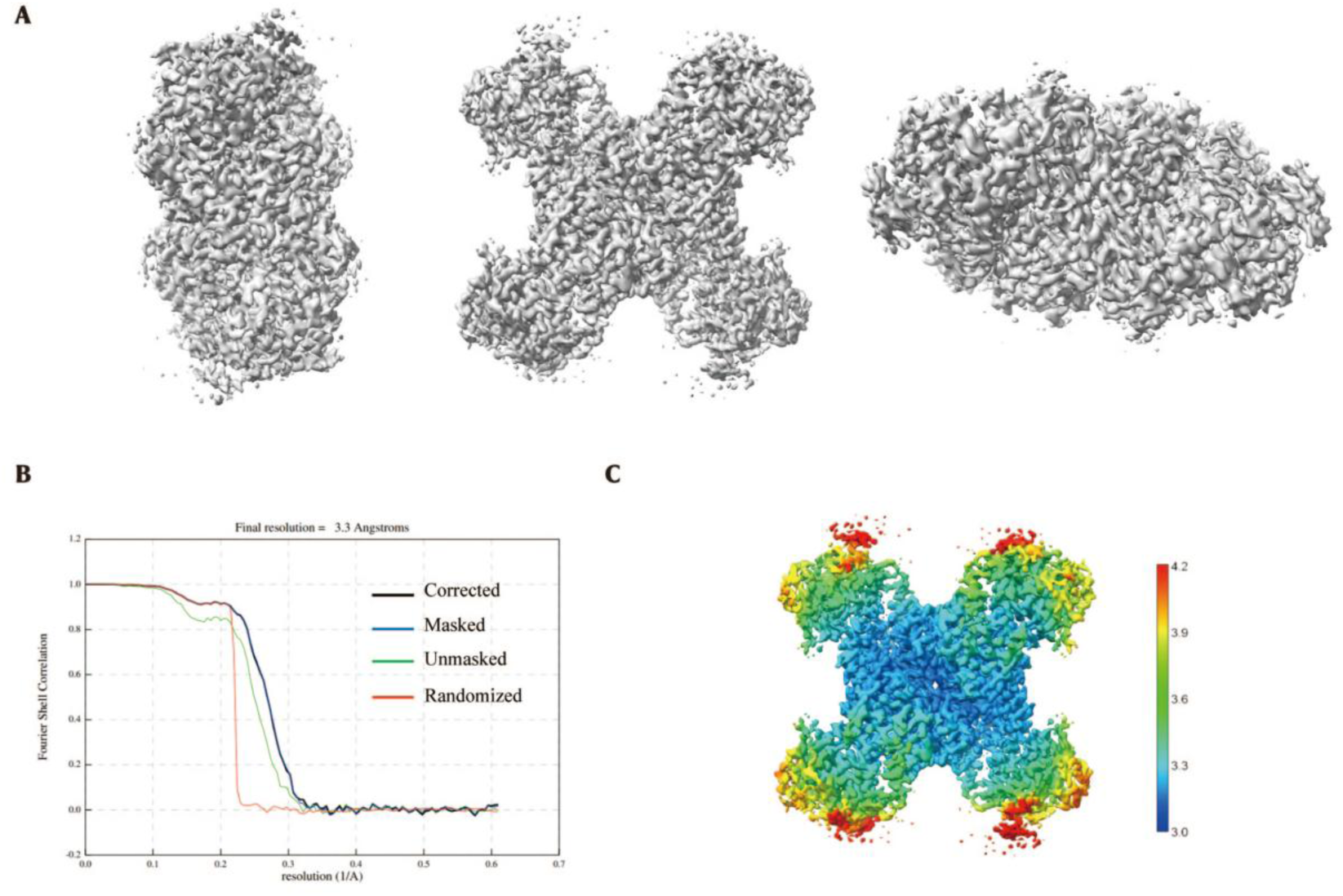
Statistics of the final density maps of the hCTPS1 p-state tetramer. (A) Three views of the final tetramer model. (B) Gold-standard FSC curve of the final density maps of the hCTPS1 bound with CTP model. (C) Local resolution map of the hCTPS1 p-state tetramer 3D refinement density map.

**Figure S3.**
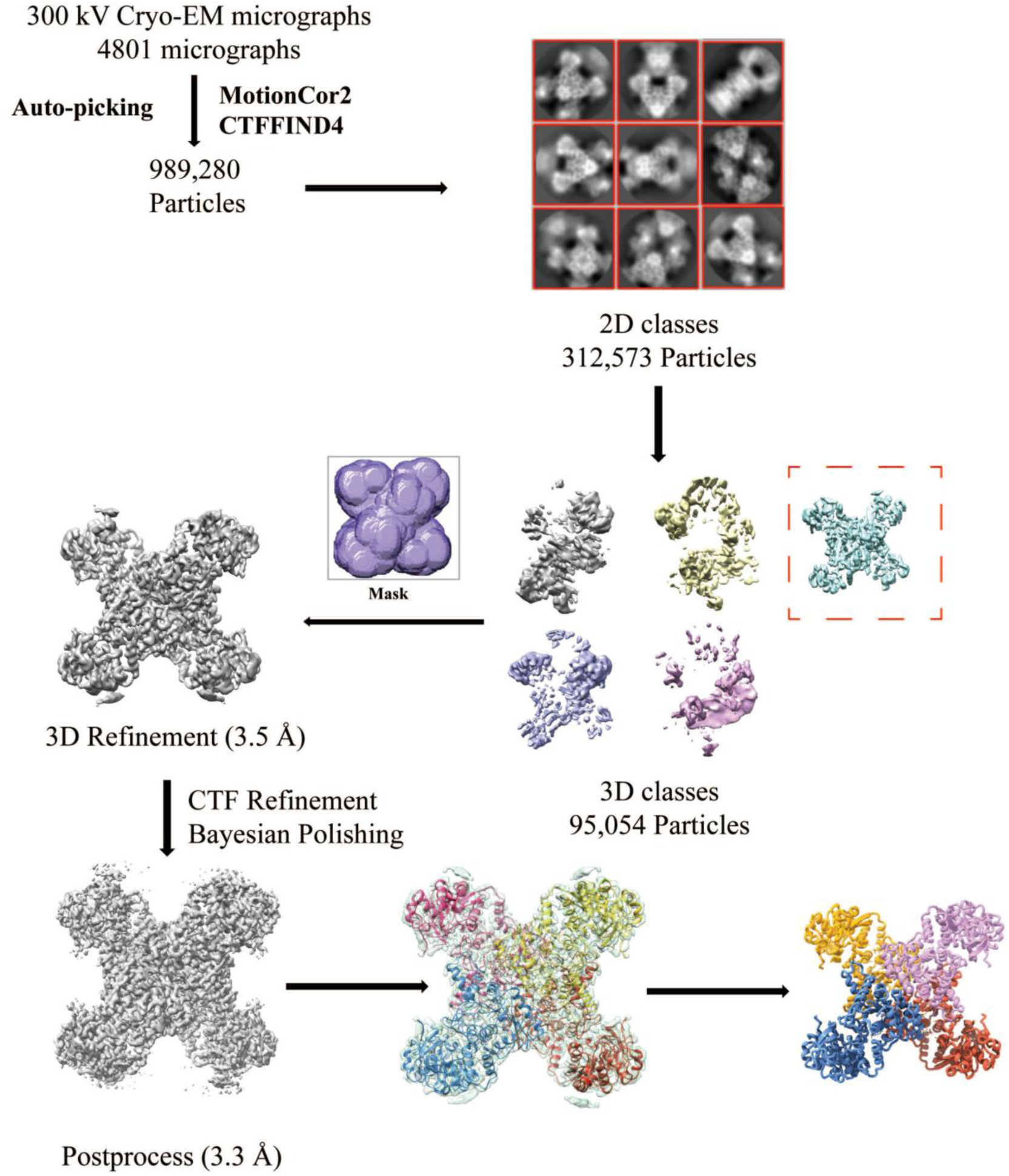
Image processing work flow of CTP-binding hCTPS1 tetramer. We collected 989,280 particles from 4801 micrographs and kept 312,573 particles after 2D classification. After 3D classification, 3D refinement and correction, final map and model was generated through different reconstructions.

**Figure S4.**
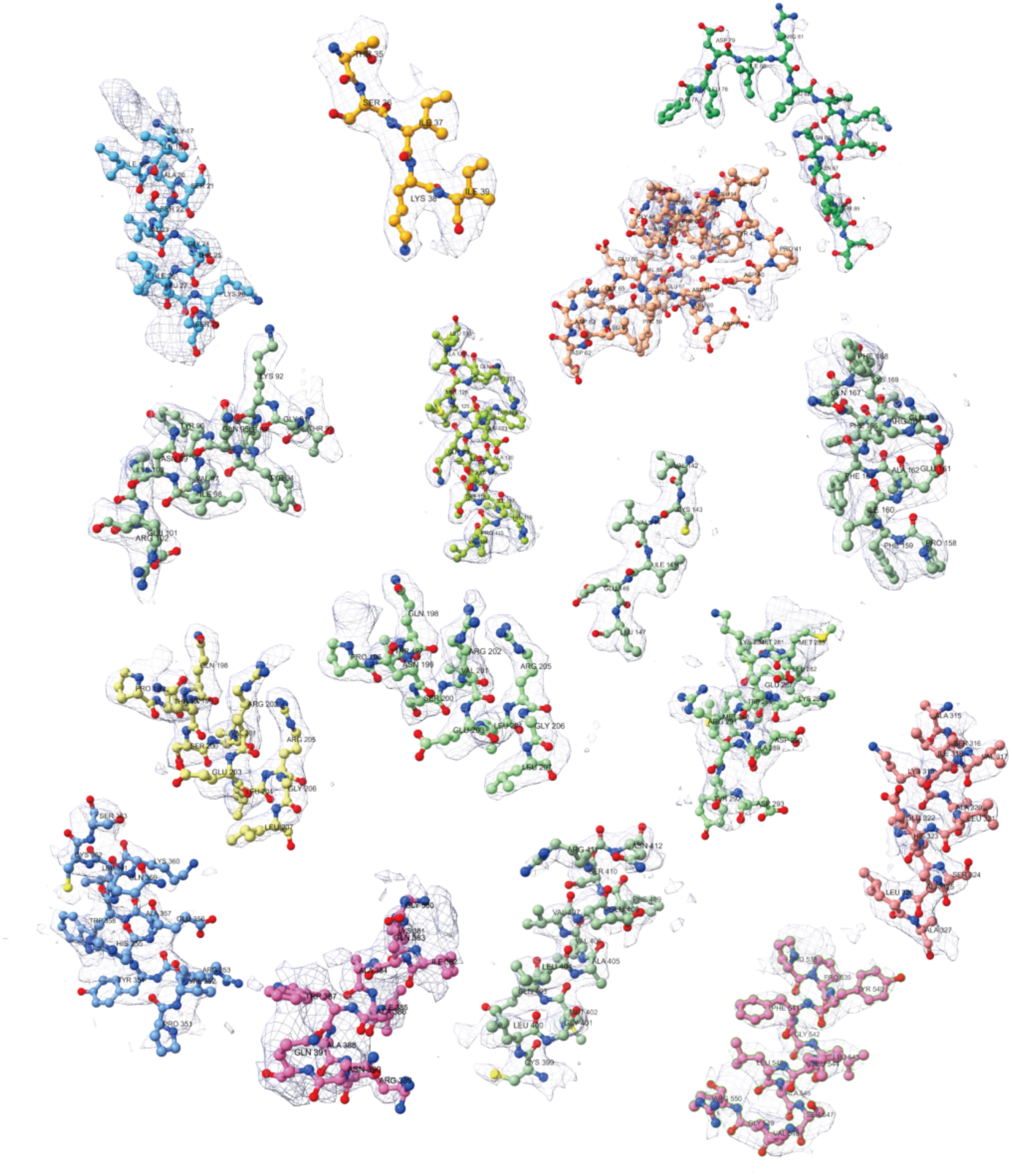
**Representative electron density map of individual regions of hCTPS1 model.**

**Figure S5.**
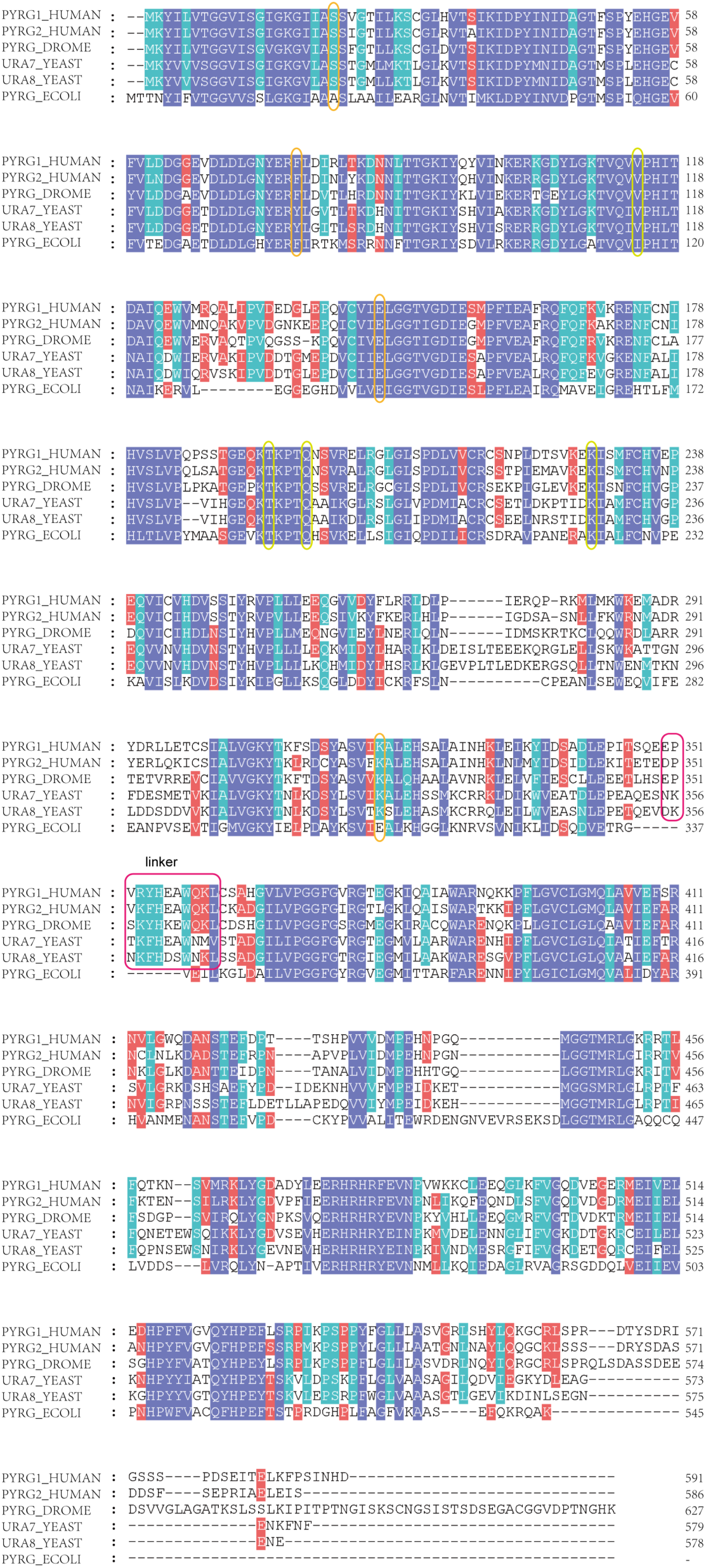
Align of CTPS sequences of various species. The boxes marked by the red represent the residue between the two domains where linker is located. The yellow boxes marked the residues involved in identification of the canonical binding pocket and the orange ones marked key residues in the non-canonical binding pocket.

**Figure S6.**
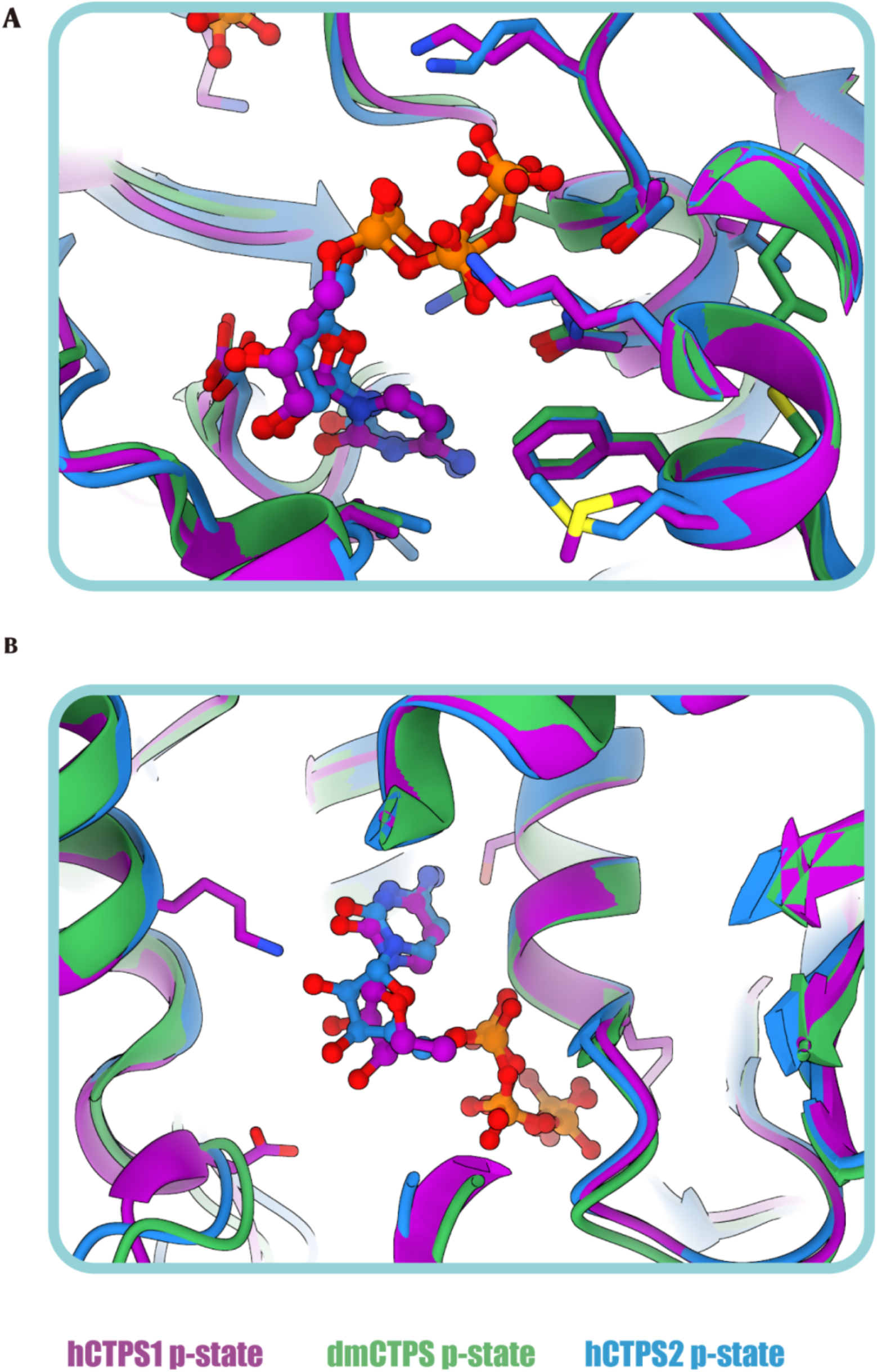
The comparison of hCTPS1, hCTPS2 and DmCTPS at two CTP binding pockets. The hCTPS1 p-state model is shown in purple, the dm-CTPS p-state model is shown in forest green and the hCTPS2 p-state model is shown in deep sky blue. (A) Canonical binding pocket (B) Non-canonical binding pocket

**Figure S7.**
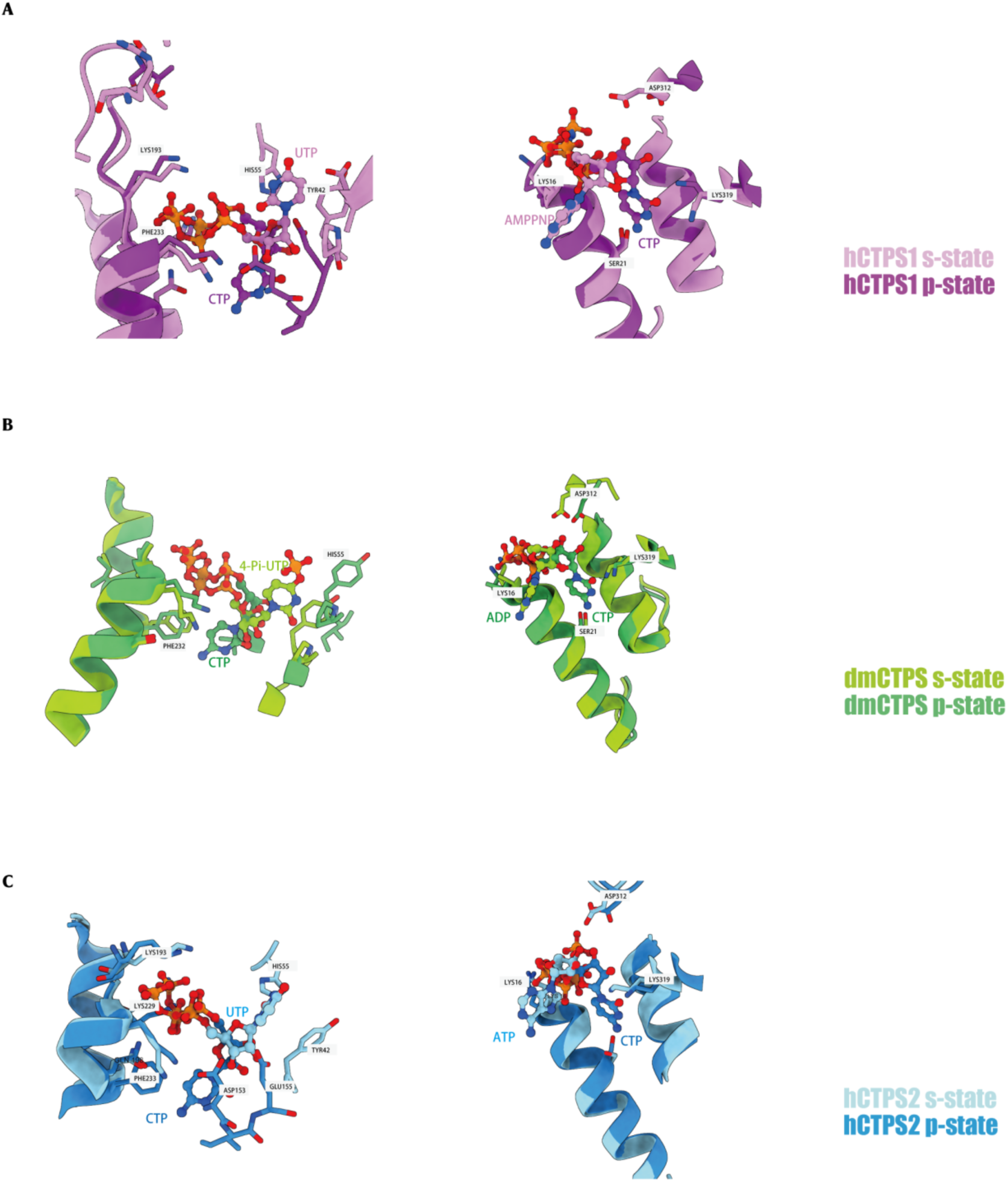
The comparison of hCTPS1, DmCTPS and hCTPS2 under substrate and product conditions. The structural of hCTPS1 (7mgz, bound with UTP and AMPPNP), DmCTPS (7dpt, bound with 4-Pi-UTP and ADP) and hCTPS2(6pk4, bound with UTP and ATP) and DmCTPS in s-state are colored in pink, lime green and powder blue, respectively. The structural of hCTPS1(9lyx, bound with CTP), DmCTPS (7dpw, bound with CTP) and hCTPS2 (7mh1, bound with CTP) in p-state are colored in purple, forest green and deep sky blue, respectively.

**Figure S8.**
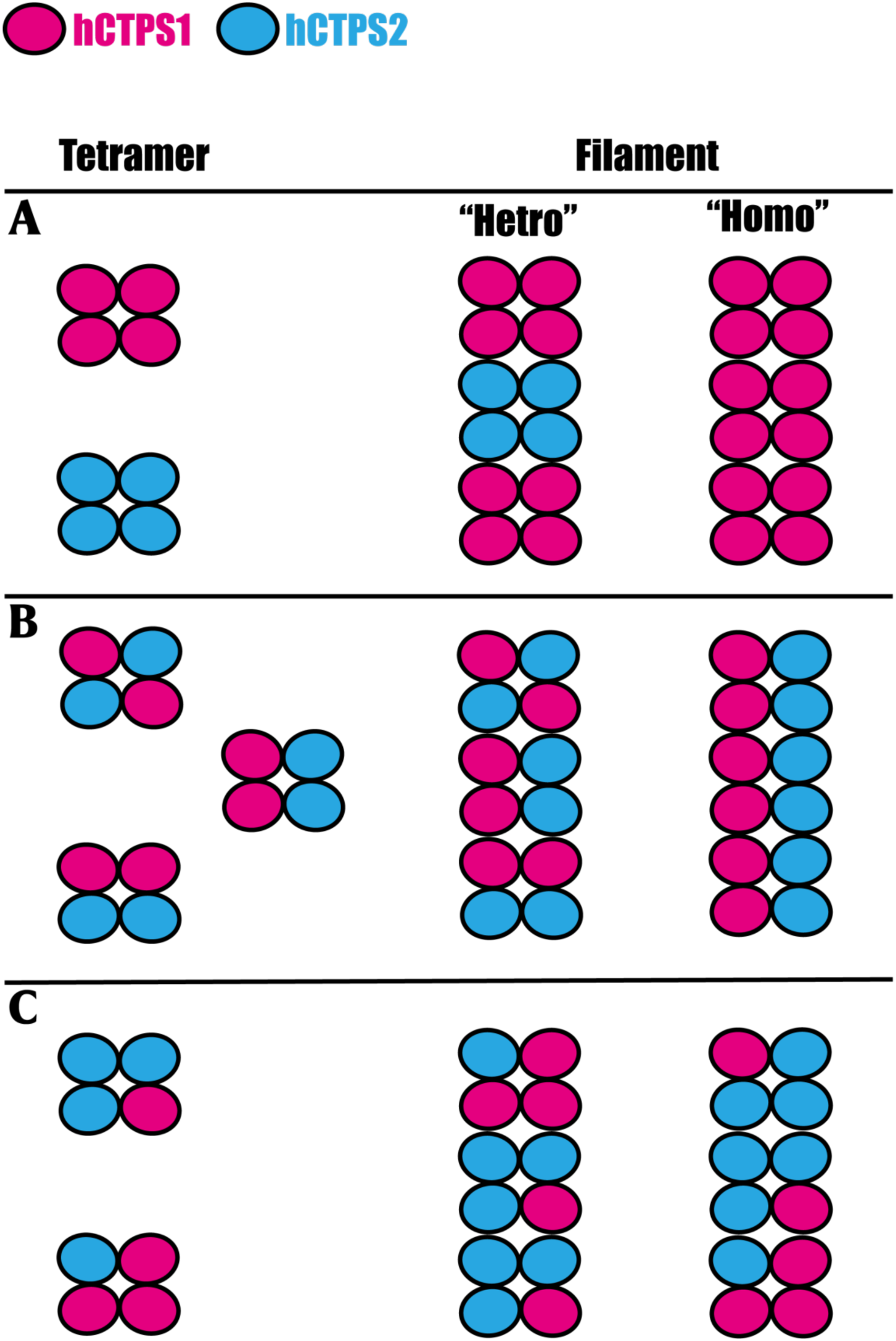
Possible assemble mechanisms of hCTPS isoforms tetramer and filament. (A) One or two homo-tetramers assemble into heterogeneous filament structures through analogous interaction interface. (B) Chimeric tetramers, each formed by 2 hCTPS1 monomers and 2 hCTPS2 monomers, with symmetrical structure, assemble into filament through binding to ligands. (C) Various hCTPS isoform monomers from heterogeneous tetramers and further assemble into irregular filament through similar interface.

---

**Table. S1.** Cryo-EM data collection and model refinement.

|  |  |
| --- | --- |
| Data collection |  |
| EM equipment | Titan Krios |
| Detector | K3 camera |
| Magnification | 29,000× |
| Voltage (kV) | 300 |
| Electron exposure ((e<sup>-</sup>/Å<sup>2</sup>)) | 60 |
| Defocus range(μm) | -1.5 to -2.5 |
| Pixel size(Å) | 0.82 |
| Symmetry imposed | D2 |
| Number of collected movies | 5387 |
| Initial particle images (no.) | 989280 |
| Final particle images (no.) | 95054 |
| Map resolution (Å) | 3.3 |
| FSC threshold | 0.143 |
| Map resolution range (Å) | 3.2-4.3 |
| Refinement |  |
| Initial model used (PDB code) | Alphafold |
| Map sharpening B-factor(Å<sup>2</sup>) | -120 |
| Model composition |  |
| Non-hydrogen atoms | 17176 |
| Protein residues | 2160 |
| Ligands | CTP |
| Water | 0 |
| Ions | 8 |
| B factors(Å<sup>2</sup>) |  |
| Protein | 54 |
| Ligand | 25 |
| Water | - |
| R.m.s. deviations |  |
| Bond lengths (Å) | 0.25 |
| Bond angles (°) | 0.50 |
| Validation |  |
| MolProbity score | 2.09 |
| Clashscore | 11.62 |
| Poor rotamers (%) | 0.68 |
| Ramachandran plot |  |
| Favored (%) | 91.18 |
| Allowed (%) | 8.44 |
| Disallowed (%) | 0.37 |

